# Differential role of prefrontal and parietal cortices in controlling level of consciousness

**DOI:** 10.1101/262360

**Authors:** Dinesh Pal, Jon Dean, Tiecheng Liu, Christopher Watson, Anthony G. Hudetz, George A. Mashour

## Abstract

There is current controversy regarding the role of prefrontal versus posterior cortices in consciousness. Clinical and correlative data have been used both to support and refute a causal role for prefrontal cortex in the level of consciousness, but a definitive relationship has not been demonstrated. We used anesthetic-induced unconsciousness as a model system to study the effect of cholinergic and noradrenergic stimulation of rat prefrontal and posterior parietal cortices on the level of consciousness. We demonstrate that cholinergic stimulation of prefrontal cortex, but not parietal cortical areas, restored wakefulness in rats despite continuous exposure to sevoflurane anesthesia. Noradrenergic stimulation of the prefrontal or parietal areas did not reverse the anesthetized state. We conclude that cholinergic mechanisms in prefrontal cortex can control the level of consciousness.

**One Sentence Summary:** Prefrontal cholinergic stimulation restores consciousness in rats despite continuous exposure to sevoflurane anesthesia

## Main Text

Activation of subcortical areas, including components of the reticular activating system, has been causally linked to behavioral arousal and levels of consciousness (*1*–*8*). By contrast, despite having been extensively investigated as a neuroanatomical substrate for the contents of consciousness, cortical control of the levels of consciousness is not well understood. Recent correlative evidence has suggested that the prelimbic prefrontal cortex might be important in regulating level of consciousness *(9)* but definitive evidence, and a comparison with more posterior cortical sites, is lacking. Addressing this question is important given the ongoing debate as to whether prefrontal or posterior cortical areas are critical for consciousness (*10*–*18*). Unlike more posterior areas of the cortex, the prefrontal cortex has extensive reciprocal connections to arousal centers in the brainstem and diencephalon, and hence is in a unique position to modulate level of consciousness (*19*,*20*). Therefore, to test the hypothesis that prefrontal cortex plays a critical role in consciousness, we attempted to reverse ongoing pharmacological coma – induced via the general anesthetic sevoflurane – by cholinergic and noradrenergic stimulation of the prelimbic prefrontal cortex and two areas in the parietal cortex.

Rats were continuously anesthetized with clinically-relevant concentrations of sevoflurane (1.9–2.4%) to maintain a constant loss of righting reflex along with complete immobility, which were used as surrogates for unconsciousness. Cholinergic stimulation was achieved via reverse dialysis of 5 mM carbachol, a mixed cholinergic agonist; noradrenergic stimulation was achieved via reverse dialysis of 20 mM noradrenaline. The concentration of carbachol was based on a previous microdialysis study in prefrontal cortex *(21)* and dose-response experiments (n=5 rats) conducted in our laboratory in which we titrated the minimum concentration of carbachol (2.5–5 mM) that produced wakefulness (data not included in this study). The concentration of noradrenaline was based on a previous dose-response study from our laboratory in which infusion of a similar concentration of noradrenaline into basal forebrain produced electroencephalographic activation and microarousals in rats under desflurane anesthesia *(4)*. We analyzed the effect of carbachol and noradrenaline delivery into the prelimbic region of prefrontal cortex and two areas in parietal cortex (posterior parietal cortex and medial parietal association cortex) on behavioral arousal, electroencephalographic activation, respiration rate, heart rate, and local acetylcholine levels (see **Fig. S1A** for experimental design).

Reverse dialysis delivery of carbachol into prefrontal cortex (n=11) during sevoflurane anesthesia induced signs of wakefulness in all 11 rats and four out of the 11 rats regained complete mobility while continuously breathing clinically relevant concentrations of sevoflurane anesthesia (1.9–2.4%) (**Fig. 1A**, see movie **S1**). The transition to wakefulness was preceded by electroencephalographic activation (**Fig. 1C**), increase in respiration rate (p=0.002) (**Fig. 1E**), and a small but significant elevation of heart rate (p=0.04) (**Fig. 1F**). Of note, carbachol-induced wakefulness was not a discrete single time event but persisted despite continuous anesthetic exposure. As opposed to cholinergic stimulation, reverse dialysis delivery of noradrenaline (n=11) into prefrontal cortex of sevoflurane-anesthetized rats did not produce any signs of wakefulness or behavioral arousal (**Fig. 1B**, see movie **S2**) but did produce electroencephalographic activation (**Fig. 1D)** and increase in respiration rate (p=0.001) (**Fig. 1G)**; there was no change in the heart rate (p= 0.2) (**Fig. 1H**). These data demonstrate that cholinergic stimulation of prefrontal cortex in the anesthetized rat is sufficient to restore wakefulness and reverse the anesthetized state.

**Figure 1:**
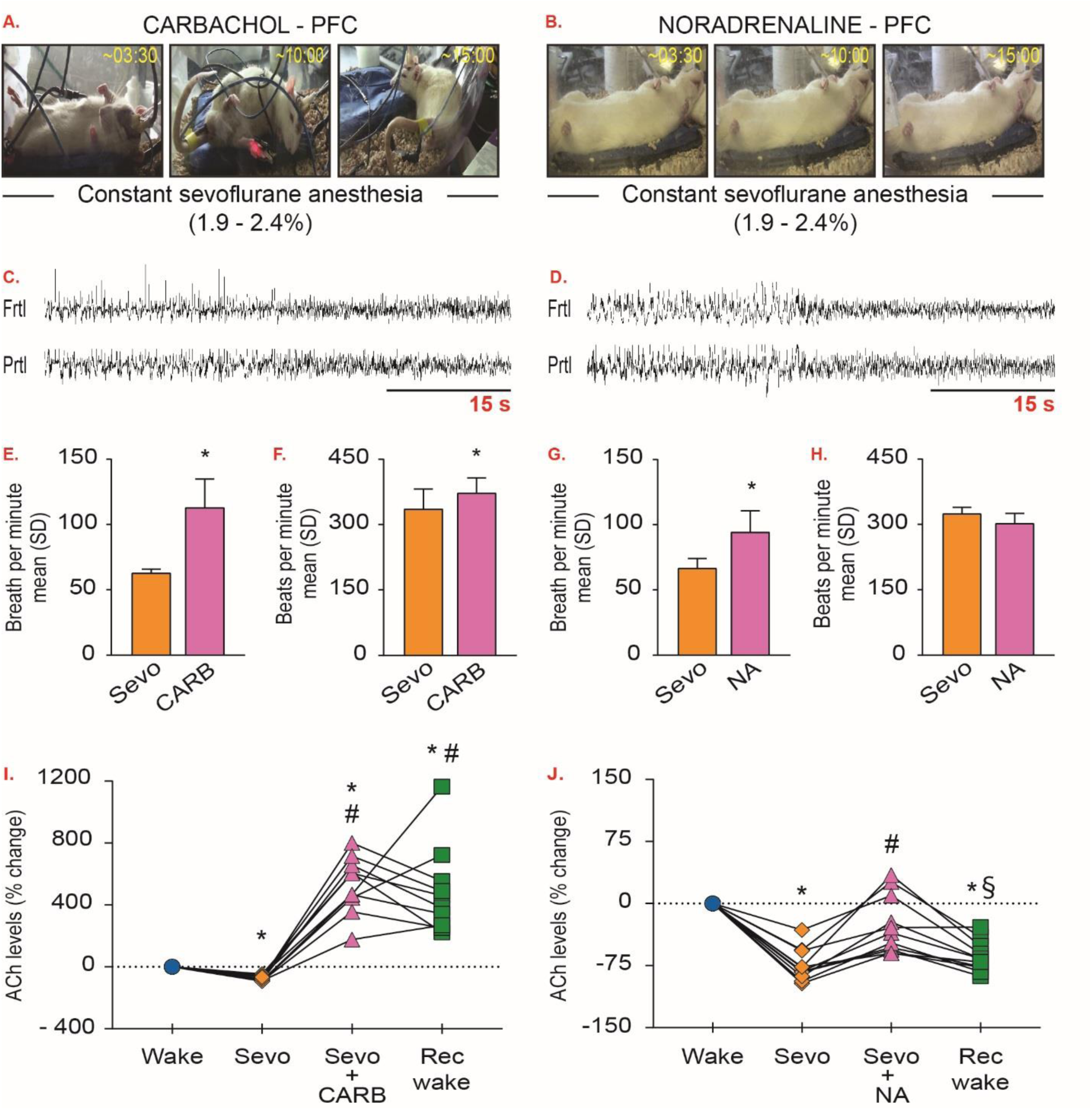
Cholinergic stimulation of prelimbic prefrontal cortex restores wakefulness in anesthetized rats. Panels in (A) and (B) show behavior after dialysis delivery of 5 mM carbachol (CARB) or 20 mM noradrenaline (NA) into prelimbic prefrontal cortex (PFC) of rats exposed to sevoflurane anesthesia (1.9–2.4%); numbers on the top right corner show the time elapsed (min:s) after CARB/NA reached the target site. (C) and (D) demonstrate electroencephalographic activation after CARB and NA, respectively. (E) and (F) shows the effect of CARB on respiration and heart rate, respectively. (G) and (H) shows the effect of NA on respiration and heart rate, respectively. (I) and (J) show changes in acetylcholine levels for each rat during preanesthesia baseline wake (Wake), sevoflurane-induced unconsciousness (Sevo), CARB/NA delivery into PFC during Sevo, and post-Sevo recovery wake (Rec wake) epochs; significance symbols show group level comparisons. Statistical comparisons were done using Student’s two-tailed paired t-test; multiple comparisons were Bonferroni corrected. Significant differences are shown at p<0.05 while the actual p values are reported in the main text. *significant compared to Wake, #significant compared to Sevo, §significant compared to CARB/NA, Frtl – frontal, Prtl – parietal, SD – standard deviation.

Consistent with our recent report (*22*), sevoflurane anesthesia produced a significant decrease in acetylcholine levels in prefrontal cortex (p<0.0001), which was reversed by local delivery of carbachol or noradrenaline (**Fig. 1I**–**J**). Carbachol in prefrontal cortex produced a substantial increase (~600%) in local acetylcholine levels as compared to both waking (p<0.0001) and sevoflurane anesthesia (p<0.0001) (**Fig. 1I**). Following noradrenaline delivery, the acetylcholine levels showed a modest but statistically significant increase as compared to sevoflurane anesthesia (~50%, p=0.001) that was not significantly different from the baseline waking state (p=0.2) (**Fig. 1 J**). The acetylcholine levels in the carbachol group stayed significantly high during the post-sevoflurane recovery epoch as compared to both waking (p=0.003) and sevoflurane anesthesia (p=0.001) epochs (**Fig. 1I**); there was no significant difference in acetylcholine levels between carbachol and recovery wake epochs (p=1). In contrast, the acetylcholine levels during the post-sevoflurane recovery wake epoch in the noradrenaline group were significantly lower as compared to both waking (p<0.0001) and noradrenaline (p=0.005) epochs (**Fig. 1J**).

In order to confirm the site specificity of the effects observed in prefrontal cortex—and to test the prevailing alternate hypothesis (*10*,*12*,*13*,*15*) that posterior cortical areas, including parietal cortex, play a critical role in consciousness—we conducted similar cholinergic stimulation (5 mM carbachol) in two distinct regions within the parietal cortex of two separate groups of rats: 1) posterior parietal cortex (n=11) which is primarily a somatosensory area, and 2) medial parietal association cortex (n=8), which is linked to cognition and attention (*23*). An additional group of rats was similarly prepared for dialysis delivery of noradrenaline into posterior parietal cortex (n=11). Carbachol delivery into posterior parietal cortex during sevoflurane anesthesia did not produce behavior consistent with wakefulness and none of the rats made any attempts at regaining the righting reflex; rats did display uncoordinated spastic muscle twitches and occasional movements in whiskers, limb and tail (**Fig. 2A**, see movie **S1**). Noradrenaline delivery into posterior parietal cortex during sevoflurane anesthesia also failed to produce any signs of wakefulness or behavioral arousal (**Fig. 2B**, see movie **S2**). Similar to cholinergic and noradrenergic stimulation of prefrontal cortex, carbachol and noradrenaline delivery into posterior parietal cortex produced electroencephalographic activation (**Fig. 2C–D**) and increase in respiration rate (p=0.0008 for carbachol group, p=0.03 for noradrenaline group) (**Fig. 2E and G**); neither carbachol nor noradrenaline affected the heart rate (p=0.06 for carbachol group, p=0.7 for noradrenaline group) (**Fig. 2F** and **H**). Sevoflurane-induced unconsciousness produced a decrease in acetylcholine levels in posterior parietal cortex (p<0.0001) in both carbachol (**Fig. 2I**) and noradrenaline (**Fig. 2J**) groups. As compared to sevoflurane epoch, dialysis delivery of carbachol and noradrenaline produced a modest but significant increase in local acetylcholine levels (p=0.002 for carbachol group, p=0.008 for noradrenaline group) that reached the wake levels (p=1 for both carbachol group and noradrenaline groups) (**Fig. 2I and J**). In a separate group of 3 rats, we delivered a higher concentration of carbachol (15 mM) into posterior parietal cortex to ensure that the concentration of carbachol is not a limiting factor for the lack of behavioral effects. The higher concentration of carbachol did not produce any significantly different behavior than that observed after 5 mM carbachol (data not included in this study). The dialysis delivery of 5 mM carbachol into medial parietal association cortex also did not produce any signs of behavioral arousal (**Fig. 3A**), but similar to the effects observed in prefrontal and parietal cortices, it did produce electroencephalographic activation (**Fig. 3B**) and increase in respiration rate (p=0.004) (**Fig. 3C**); there was no statistical change in heart rate (p=0.05) (**Fig. 3D**). Sevoflurane produced a significant decrease in acetylcholine levels in medial parietal association cortex (p<0.0001), which was not reversed by the local carbachol delivery (**Fig. 3E**). There was no difference in the acetylcholine levels between wake and carbachol (p=0.2) or carbachol and recovery (p=0.3) epochs. As compared to sevoflurane epoch, the acetylcholine levels increased during the post-sevoflurane recovery epoch (p=0.007) and reached the wake levels (**Fig. 3E**).

**Figure 2:**
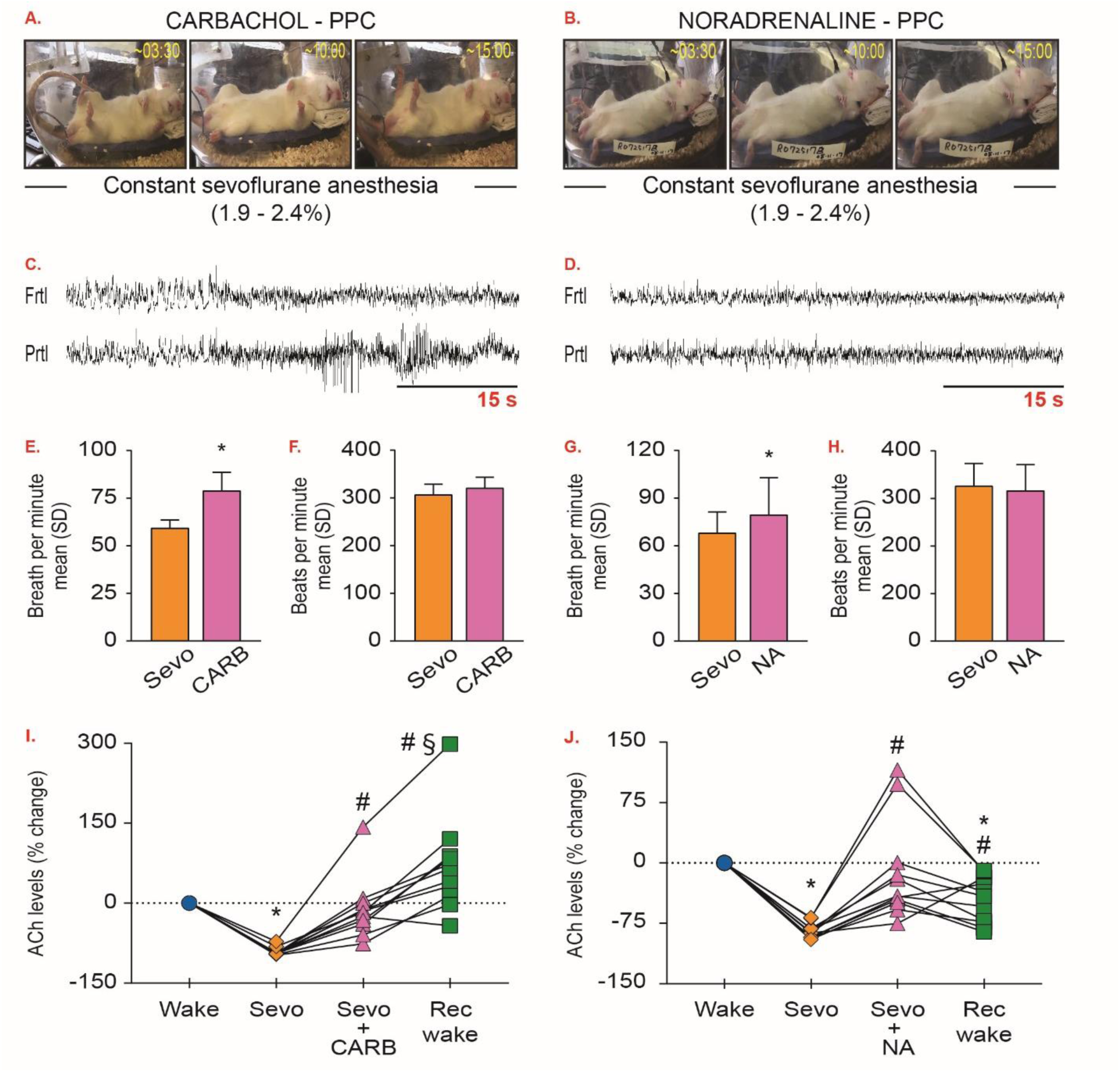
Cholinergic stimulation of posterior parietal cortex is not sufficient to restore wakefulness in anesthetized rats. Panels in (A) and (B) show behavior after dialysis delivery of 5 mM carbachol (CARB) or 20 mM noradrenaline (NA) into posterior parietal cortex (PPC) of rats exposed to sevoflurane anesthesia (1.9–2.4%); numbers on the top right corner show the time elapsed (min:s) after CARB/NA reached the target site. (C) and (D) demonstrate electroencephalographic activation after CARB and NA, respectively. (E) and (F) shows the effect of CARB on respiration and heart rate, respectively. (G) and (H) shows the effect of NA on respiration and heart rate, respectively. (I) and (J) show changes in acetylcholine levels for each rat during preanesthesia baseline wake (Wake), sevoflurane-induced unconsciousness (Sevo), CARB/NA delivery into PPC during Sevo, and post-Sevo recovery wake (Rec wake) epochs; significance symbols show group level comparisons. Statistical comparisons were done using Student’s two-tailed paired t-test; multiple comparisons were Bonferroni corrected. Significant differences are shown at p<0.05 while the actual p values are reported in the main text. *significant compared to Wake, #significant compared to Sevo, §significant compared to CARB/NA, Frtl – frontal, Prtl – parietal, SD – standard deviation.

**Figure 3:**
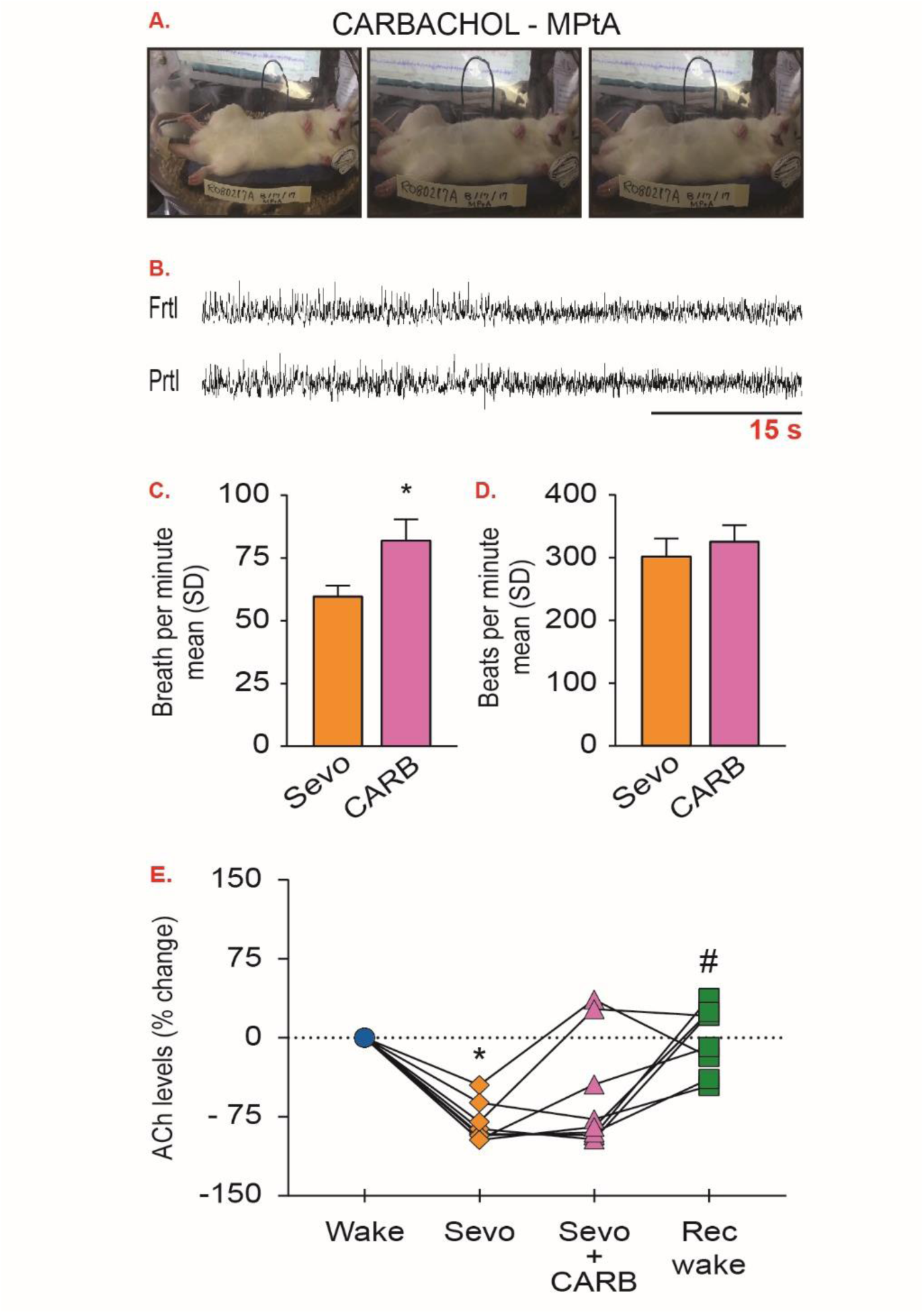
Cholinergic stimulation of medial parietal association cortex is not sufficient to restore wakefulness in anesthetized rats. A) Effect on behavior after dialysis delivery of 5 mM carbachol (CARB) into medial parietal association cortex (MPtA) of rats exposed to sevoflurane anesthesia (1.9–2.4%). B) representative traces show electroencephalographic activation after CARB delivery into MPtA. C) and (D) shows the effect of CARB delivery on respiration and heart rate, respectively. E) shows changes in acetylcholine levels for each rat during preanesthesia baseline wake (Wake), sevoflurane-induced unconsciousness (Sevo), CARB delivery into MPtA during Sevo, and post-Sevo recovery wake (Rec wake) epochs; significance symbols show group level comparisons. Statistical comparisons were done using Student’s two-tailed paired t-test; multiple comparisons were Bonferroni corrected. Significant differences are shown at p<0.05 while the actual p values are reported in the main text. *significant compared to Wake, #significant compared to Sevo, Frtl – frontal, Prtl – parietal, SD – standard deviation.

The post-sevoflurane recovery period in the rats that received carbachol into prefrontal cortex was characterized by generalized behavioral seizures and epileptiform electroencephalographic pattern. In contrast, carbachol delivery into either posterior parietal cortex or medial parietal association cortex did not produce any signs of behavioral seizures or abnormal electroencephalographic patterns during post-sevoflurane recovery epoch. However, 3/11 rats in the posterior parietal group showed intermittent spike-waveform pattern in the electroencephalogram immediately after carbachol delivery (**Fig. 2C**). Noradrenaline delivery into either prefrontal or posterior parietal cortex did not produce any abnormal electroencephalographic pattern or behavioral seizures in any of the epochs.

These data demonstrate that while cholinergic and noradrenergic stimulation of prefrontal and parietal cortices can activate the cortex, only cholinergic stimulation of the prefrontal cortex restored level of consciousness and reversed the anesthetized state. These findings are also supported by recent work in human patients demonstrating a positive effect of prefrontal stimulation on arousal in patients with pathologic disorders of consciousness *(14)*, and a longstanding hypothesis that acetylcholine is a neurochemical correlate of the capacity for consciousness (*24*,*25*). Furthermore, a recent electrophysiological study postulated that the prelimbic prefrontal cortex in rat, as was targeted in the current study, is a neural node in thalamocortical—and likely corticocortical—interactions that modulates both the induction of and emergence from propofol anesthesia *(9)*. It was unexpected that noradrenaline failed to induce any behavioral change despite its known role in arousal (*3*,*4*,*6*). This finding could relate to the pharmacokinetics of noradrenaline and carbachol. However, a more likely explanation is offered by a recent study in rat barrel cortex in which cholinergic stimulation through carbachol infusion during urethane anesthesia produced a tonic increase in firing rates whereas infusion of noradrenaline suppressed the overall firing rate *(26)*. Nevertheless, similar to carbachol infusion, noradrenaline did produce electroencephalographic activation and increase in respiration rate, thereby demonstrating that the cortical dynamics can be dissociated from behavior, i.e., an activated electroencephalogram in the absence of wakefulness. Similar dissociation between delta power—an electroencephalographic correlate of quiescence—and behavior has been reported during sleep restriction, immune challenges, dietary changes, aging, and anesthesia (*27*,*28*), which urges caution in the singular reliance on electroencephalogram-based measures as a signature of consciousness.

Our study establishes the sufficiency of cholinergic stimulation in prefrontal cortex for inducing wakefulness and restoring the level of consciousness. However, it does not exclude the possibility that the recruitment of additional brain areas, such as the posterior cortical “hot zone” proposed to be important for conscious contents (*10*,*12*,*13*,*15*), could be required to restore the complete spectrum of consciousness. Sevoflurane is known to inhibit both nicotinic and muscarinic receptors *(29*,*30*) but based on our study we cannot distinguish between the relative contributions of muscarinic and nicotinic receptors in mediating carbachol-induced reversal of sevoflurane anesthesia. In addition, even though carbachol-induced increase in prefrontal acetylcholine and the simultaneous behavioral arousal suggests a causal link between the levels of cortical acetylcholine and behavioral arousal, our study cannot rule out the possibility that increase in cortical acetylcholine levels is an epiphenomenon of behavioral arousal. Finally, we can only interpret these data in terms of the objectively observable level of consciousness (e.g., signs of wakefulness) and cannot comment definitively on how prefrontal versus posterior cortical areas contribute to the phenomenal contents of consciousness. However, it is becoming more widely appreciated that levels and contents of consciousness cannot be completely dissociated *(31)* so these findings might ultimately be relevant to the ongoing debate regarding the cortical sites that are critical for conscious experience.

In summary, these findings suggest that the prefrontal cortex is capable of regulating levels of consciousness and reversing the anesthetized state. The homology between rodent and human prefrontal cortex, and the well-characterized tripartite circuitry between cholinergic basal forebrain and the prefrontal and parietal cortices *(19*,*20)*, encourages further work to develop a more precise understanding of the mechanism by which the prefrontal cortex regulates consciousness *(16*,*17*,*18)*.

## Acknowledgments

The authors would like to thank Chris Andrews of the University of Michigan Center for Statistical Consultation and Research unit for help with statistical analysis, and Donald C. Fedrigon III for help with data collection. This work was funded by the National Institutes of Health (Bethesda, MD, USA) (R01GM098578 to GAM), and funding from the Department of Anesthesiology, University of Michigan Medical School, Ann Arbor. DP and GAM designed the study, interpreted the data, and prepared the manuscript; JD and AGH contributed to data interpretation and manuscript preparation; DP, JD, and TL conducted the experiments and collected the data; DP, JD, TL, and CW analyzed the data. All authors read and approved the manuscript. The authors declare no competing interests. All data are available in the main text or the supplementary materials. Videos showing behavior of some of the rats for each experimental groups can be made available on request to the corresponding author.

